# Pocket similarity-guided selection of protein targets for systematic protein–ligand dataset generation

**DOI:** 10.64898/2026.11.10.765562

**Authors:** Montserrat García-Cerqueda, Matthieu Schapira, Albert A Antolin, Aled M Edwards, Patrick Aloy

## Abstract

The scarcity of large, diverse, and unbiased protein–ligand binding datasets limits the performance of machine learning approaches in drug discovery, particularly for understudied proteins. The Target 2035 initiative aims to address this gap by generating open-access binding data for more than 2,000 human proteins, a goal whose value depends on maximizing the diversity of the selected targets. Because proteins with different folds can share chemically analogous binding sites, we reasoned that pocket similarity, rather than sequence or fold similarity, is the appropriate criterion for selecting representative targets that maximize coverage of the human pocketome. Here, we use PocketVec, an alignment-free descriptor of binding sites, to map the human globular pocketome, identifying 51,535 predicted druggable pockets across 15,559 proteins. A two-level hierarchical clustering, first within protein families and then across the whole proteome, grouped these pockets into 1,865 clusters of chemically similar binding sites. After discarding 533 clusters containing pockets with known ligands, we selected the 100 most populated of the remaining 1,332 ligand-free clusters and chose two high-quality representative pockets per cluster, ranked by their resemblance to known ligand-binding sites. Finally, we implemented two complementary prioritization strategies, one favoring structurally unexplored proteins and the other proteins amenable to experimental follow-up, and selected 200 pockets from 100 different clusters from each strategy, with 89 clusters in common. Both selections cover over 10,000 pockets in ∼5,600 proteins, approximately 36% of the human proteins with annotated druggable pockets, which represents a 2.8-to 4.4-fold increase in the number of covered pockets, domains, and proteins than random selection baselines. This pocket-based framework offers a principled strategy to select diverse targets for Target 2035 and other large-scale protein–ligand data generation efforts in chemical biology.

## Introduction

Despite decades of investment, only a small fraction of the approximately 20,000 proteins encoded by the human genome has been explored as therapeutic targets. Research attention follows a strong power-law distribution—a handful of well-studied proteins account for the vast majority of published studies—leaving a large portion of the proteome functionally uncharacterized and pharmacologically untouched. This so-called “dark proteome” is not a scientific backwater: it includes many proteins with compelling disease relevance whose study has been hampered by the lack of chemical tools to perturb their function^1,2^.

Against this backdrop, computational approaches have emerged as powerful allies in drug discovery. Virtual screening, machine learning (ML), and artificial intelligence (AI) methods can, in principle, rapidly traverse the enormous chemical space of potential drugs and identify promising candidates for experimental follow-up^3^. Recent years have seen a proliferation of AI-based platforms for hit finding, *de novo* drug design, and target identification^4–6^. Yet the practical impact of these methods remains constrained by a fundamental bottleneck: the availability of sufficiently large, diverse, and unbiased protein–ligand binding datasets on which to train and validate predictive models^7,8^. Most existing datasets are heavily skewed toward a small number of well-studied protein families—kinases, GPCRs, proteases—reflecting historical research priorities rather than biological diversity. Indeed, the Critical Assessment of Computational Hit-finding Experiments (CACHE) are dynamically monitoring progress and, more importantly, highlighting the limitations of current methods and training datasets^9^.

Central to structure-based drug discovery is the concept of the druggable pocket: a protein cavity capable of binding an organic small molecule with sufficient affinity and selectivity to modulate the protein’s function. Detecting and characterizing such pockets at proteome scale is therefore a prerequisite for any systematic chemical biology effort. Pocket detection algorithms can be broadly grouped into two classes: ligand-based approaches, which define binding sites from co-crystallized ligands observed in experimentally determined structures, and *de novo* detection algorithms, which identify geometrically and physicochemically favorable cavities in any structure—experimental or predicted. Among the latter, Fpocket^10^ uses a Voronoi tessellation approach to rapidly enumerate cavities, while P2Rank^11^ applies random-forest classification trained on local surface properties to rank predicted pockets by druggability. The parallel use of both ligand-based and *de novo* approaches, applied to both experimentally determined and AlphaFold2 (AF2) predicted structures, offers the broadest possible coverage of the human pocketome^12^.

Once detected, binding sites must be characterized in a computationally tractable format that enables the comparison of any two pockets, regardless of protein family or fold. Existing pocket descriptor strategies fall into several broad categories^13,14^. Residue-based methods (e.g. FuzCav^15^, SiteAlign^16^) encode pockets from the physicochemical properties of their constituent amino acids. Surface-based methods (e.g. MaSIF^17^) represent the molecular surface of the binding site. Interaction-based methods (e.g. KRIPO^18^, TIFP^19^) explicitly encode protein– ligand contacts, but require co-crystallized ligands and are therefore limited to *holo* structures. Data-driven methods (e.g. DeeplyTough^20^) learn pocket representations from structural data using deep learning, but often at the cost of interpretability. Despite their diversity, all these approaches share a common limitation: their descriptors are hand-crafted or trained on specific benchmark datasets and can generalize poorly to novel protein families or predicted structural models. Moreover, alignment-dependent methods incur prohibitive computational costs at proteome scale.

Recently, we presented PocketVec, a new strategy based on inverse virtual screening designed to overcome these limitations^14^. Rather than directly encoding the geometry or chemistry of a pocket, it exploits a refined form of the chemogenomics principle—*similar pockets bind similar ligands*—to generate fixed-length, alignment-free descriptors via inverse virtual screening. A standardized library of 128 lead-like molecules is docked against each pocket using rDock rigid docking, and the resulting docking scores are converted into rankings that form the descriptor vector. Any two pockets can then be readily compared by computing the cosine distance between their descriptors, making proteome-wide all-against-all comparisons tractable. Benchmarked on the ProSPECCTs dataset^13^, PocketVec achieves an AUROC of 0.97 on the primary orthosteric-pair benchmark set, and validation on over two million protein pairs from ChEMBL and BindingDB confirms that lower cosine distance reliably predicts a higher rate of shared bioactive compounds. Importantly, PocketVec operates on any three-dimensional structure—experimental or AF2-predicted—and does not require co-crystallized ligands, making it well suited to characterize the full human pocketome including the majority of proteins for which no inhibitor is yet known. Indeed, we were able to identify and characterize over 30,000 potentially druggable pockets in human proteins.

The Target 2035 initiative, led by the Structural Genomics Consortium (SGC), was founded to break the data bottleneck that constrains AI-based drug discovery^1,2^. Its current phase (2025–2030) aims to systematically generate open-access, high-quality protein–small-molecule binding datasets for more than 2,000 human proteins using affinity-selection mass spectrometry (AS-MS) and DNA-encoded chemical library (DEL) screening, making all data freely available through the AIRCHECK database^21,22^. The scientific value of this effort depends critically on maximizing the diversity of the selected targets: as the Target 2035 roadmap explicitly states, “the selection of which proteins to screen will be guided by the requirement to maximize the structural and functional diversity of the protein targets”^21^.

The challenge of selecting a maximally diverse and representative subset of proteins is reminiscent of earlier structural genomics initiatives, which resolved it by clustering proteins by structural similarity and selecting one representative per family^23–25^. We can borrow this logic for Target 2035, but the appropriate grouping criterion must shift from fold similarity to pocket similarity: two proteins with different overall architectures but chemically equivalent binding sites will tend to bind similar compounds, so characterizing one representative provides information relevant to both. Here, we identify over 50,000 potentially druggable pockets in the globular human proteome, compute their PocketVec descriptors and use them to cluster them according to their similarity. Finally, we propose two complementary target-selection strategies to identify 200 representative pockets that maximize the coverage of the globular human pocketome.

## Results and Discussion

### An updated and expanded map of the human druggable pocketome

To obtain an updated version of the globular human druggable pocketome, we first generated an updated and expanded set of pocket descriptors covering all druggable cavities in the human proteome. Starting from the complete set of human protein sequences in UniProt^26^, we annotated protein domains using Pfam^27^ via InterPro^28^, including entries classified as either “domain” or “family”. This yielded 41,774 domains spanning 7,187 Pfam families across 17,690 proteins—a 49% increase over the 28,044 domains in the original publication^14^. The increase reflects three additions relative to the original pipeline: (i) the incorporation of Pfam annotations at the “family” level, which expand coverage to proteins lacking detailed structural annotation; (ii) the inclusion of mmCIF-format PDB entries^29^ and (iii) AF2 models for 1,007 domains from proteins exceeding 2,700 amino acids, which were excluded from the original implementation.

For structural annotation, we retrieved experimental structures with at least 80% domain coverage from the PDB^30^, yielding at least one structure for 12,572 domains (3,824 Pfam families, 7,145 proteins). AF2 structural models^31^ covered 41,565 domains (7,167 Pfam families, 17,621 proteins). For each annotated domain, we identified candidate druggable pockets using a ligand-based approach (LIG) and a *de novo* pocket discovery approach (PD), and generated PocketVec descriptors for all identified pockets using the same docking-based inverse virtual screening pipeline, with structure preparation performed by PDBFixer^32^ as an open-source replacement for the original MOE-based workflow^14^. See Figure 1a and *Methods* for details.

**Figure 1.**
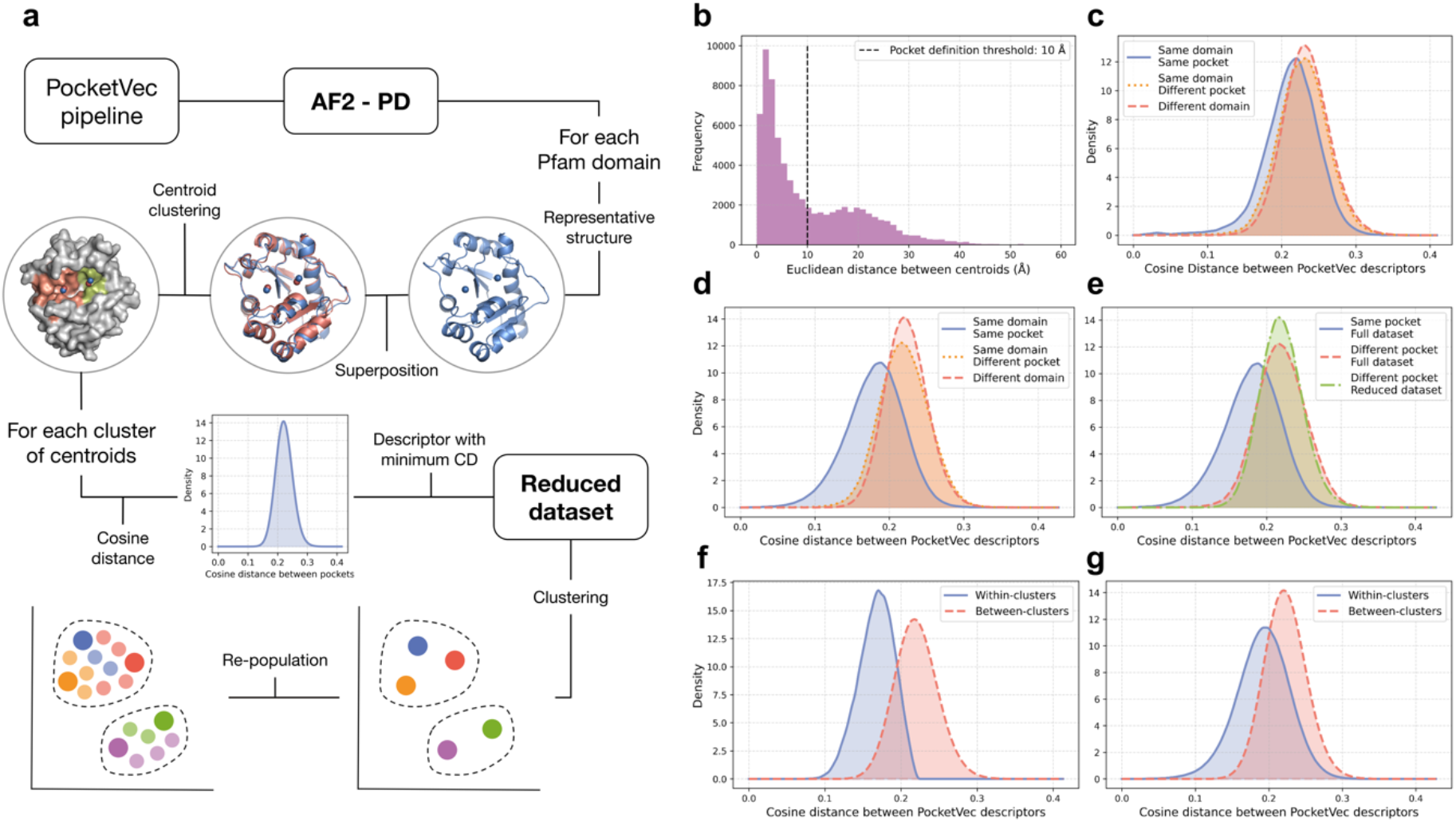
Obtaining clusters of similar ligand-binding pockets. (**a**) Overview of the two-stage clustering pipeline. For each Pfam domain in the AF2-PD dataset, a reference structure is selected and all other domain structures are superimposed; pocket centroids are clustered to obtain shared Pfam-level pockets. A representative centroid is selected per Pfam-level pocket (minimum average cosine distance), and the resulting reduced dataset is clustered by cosine distance using complete-linkage agglomerative clustering. Non-representative centroids are then re-assigned to the cluster of their representative. (**b**) Histogram of pairwise Euclidean distances between pocket centroids from the same Pfam domain in the PDB-LIG dataset; the 10 Å threshold is indicated. (**c**) Cosine distance distributions between centroids belonging to the same Pfam-level pocket (blue), different pockets within the same domain (orange), and different domains (red) for the PDB-LIG dataset. (**d**) Same as (c) for the AF2-PD dataset. (**e**) Inter-pocket cosine distance distributions for the full (red) and reduced (green) AF2-PD datasets, and intra-pocket cosine distances (blue). (**f**) Intra-(blue) and inter-(red) cluster cosine distance distributions for the reduced AF2-PD dataset. (**g**) Same as (f) after re-population of clusters with non-representative centroids.

The updated datasets represent a substantial expansion across all four pocket sets. The AF2-PD dataset, which is the most comprehensive, grew from 32,202 to 51,535 pockets—a 60% increase that adds more than 19,000 novel binding sites distributed across 11,624 additional domains and more than 5,000 additional proteins relative to the original characterization. The LIG datasets nearly doubled (PDB-LIG +91%; AF2-LIG +93%), reflecting both increased Pfam annotation coverage and the incorporation of mmCIF structures. Together, these datasets demonstrate that the updated pipeline substantially extends the reach of the original human pocketome map.

Finally, we generated the PocketVec descriptors following the procedure described in the original publication (see *Methods*). The final number of pockets, domains, Pfam domains and proteins contained in each dataset is detailed in Table 1. In the original PocketVec study, all four datasets were benchmarked against ProSPECCTs ^13^ and validated through large-scale comparison against ChEMBL and BindingDB binding data, establishing their accuracy as descriptors of pocket similarity. We repeated the ProSPECCTs benchmarking to calibrate the similarity thresholds for the updated descriptors (Supplementary Figures 1 and 2), confirming that descriptor quality is maintained across datasets. Having established the validity of all four sets, we focus the remainder of the analyses exclusively on the AF2-PD dataset, which provides the largest coverage of the human proteome— 15,559 proteins versus 7,030 for PDB-PD and fewer than 2,100 for either LIG dataset—and includes the many disease-relevant proteins for which no experimental structure or known ligand is available. The other three datasets are provided as a resource for community use.

**Table 1.** Number of potentially druggable pockets, globular domains, unique Pfam families and proteins identified and annotated from experimental structures (PDB) and AlphaFold2 models (AF2) using a ligand-based (LIG) or *de novo* pocket detection (PD) approach.

|  | PDB-LIG | PDB-PD | AF2-LIG | AF2-PD |
|---|---|---|---|---|
| N<sup>o</sup> of pockets | 3,069 | 23,326 | 2,705 | 51,535 |
| N<sup>o</sup> of domains | 2,129 | 12,081 | 1,990 | 30,835 |
| N<sup>o</sup> of Pfam | 736 | 3,738 | 698 | 5,875 |
| N<sup>o</sup> of proteins | 2,045 | 7,030 | 1,869 | 15,559 |

### Defining shared binding sites within protein families

A fundamental challenge in working with the complete pocketome is structural redundancy: multiple structural models of the same domain—from different crystal forms, resolution, or experimental conditions—generate nearly identical pocket descriptors. Propagating this redundancy into downstream clustering would inflate the apparent size of some protein families and bias representative selection.

To address this, we defined shared, family-level binding sites by grouping pockets from different structural copies of the same Pfam domain. For each Pfam family represented by at least two domain structures, we superimposed all domain models onto a single reference using TM-align^33^ and clustered pocket centroids by pairwise Euclidean distance using complete-linkage agglomerative hierarchical clustering. The distance threshold of 10 Å was determined from the observed bimodal distribution of centroid-to-centroid distances within Pfam families, which separates proximal centroids from the same pocket (first peak at ∼2–3 Å) from centroids belonging to distinct pockets of the same domain (second peak at ∼18–20 Å; Figure 1b). Each resulting cluster of centroids was treated as a single shared Pfam-level pocket.

We validated this pocket definition schema on the experimentally determined PDB-LIG dataset, where 736 Pfam families yielded 1,265 Pfam-level pockets. Cosine distance distributions confirmed that centroids within the same Pfam-level pocket (mean 0.213 ± 0.038) were statistically more similar to each other than centroids from different pockets of the same domain (0.228 ± 0.034; KS test D = 0.166, p=0) or from different domains (0.233 ± 0.031; D = 0.216, p=0; Figure 1c). We thus applied the same strategy to the 5,875 Pfam families in the AF2-PD dataset, which yielded 17,669 Pfam-level pockets, with an even clearer separation between distance distributions (same pocket: 0.18 ± 0.039; different pocket, same domain: 0.22 ± 0.033, D = 0.415, p=0; different domain: 0.223 ± 0.029, D = 0.475, p=0; Figure 1d), confirming robust pocket definition across both structure sources.

The 17,669 Pfam-level representative pockets represent a 65.7% reduction from the original 51,535 AF2-PD descriptors. We verified that the inter-descriptor distance distribution of the reduced dataset closely matched that of the full dataset (Figure 1e), ensuring that removing intra-family redundancy does not distort the global pocket landscape. Each Pfam-level pocket is represented by a single centroid: the descriptor with the minimum mean cosine distance to all other members, chosen as the most representative of the group.

### Clustering the pocket landscape across the entire proteome

Pfam-level grouping removes intra-family redundancy but does not capture similarity between pockets in different protein families. Discovering such cross-family relationships is central to the PocketVec approach, as it allows a single representative pocket to stand in for chemically related binding sites distributed across unrelated proteins.

Thus, we next grouped the 17,669 reduced AF2-PD descriptors using complete-linkage agglomerative hierarchical clustering with a cosine distance threshold of 0.22, obtaining 1,865 distinct clusters. This threshold was set at approximately the median inter-pocket distance to apply a permissive grouping criterion that compensates for the strict complete-linkage criterion (which requires all members within a cluster to remain within the threshold of each other). The resulting clusters showed a clear separation between intra-cluster (0.168 ± 0.023) and intercluster (0.22 ± 0.029) cosine distance distributions (KS test D = 0.69, p = 0; Figure 1f), confirming that pockets within the same cluster are substantially more similar to each other than to pockets in other clusters. We also explored the use of a cosine distance threshold of 0.16 in the clustering, identified as optimal to distinguish similar pockets in the ProSPECCT benchmark. Indeed, while this more stringent threshold increased the difference in pocket similarity distributions (Supplementary Figure 3a), it also resulted in smaller clusters, many contained only one pocket (Supplementary Figure 3b).

Finally, to maximize the pool of candidates available for target selection, we re-populated each cluster by assigning all non-representative Pfam-level pockets to the cluster of their corresponding representative. This slightly increased the average intra-cluster distance (0.192 ± 0.037) but the inter/intra separation remained highly significant (KS test D = 0.368; Figure 1g). The re-populated clusters collectively cover 51,535 AF2-PD pockets across 30,835 domains and 15,559 proteins—the full breadth of the dataset—each organized into a chemically coherent neighborhood of pocket space.

### Final selection of representative pockets

With 1,865 clusters available, we applied a systematic filtering procedure to focus on clusters representing novel, chemically uncharacterized pocket space (Figure 2a). Since the goal of Target 2035 is to generate binding data for proteins without known modulators, we excluded clusters containing at least one pocket with a co-crystallized ligand. Ligand association was established by superimposing AF2-LIG and AF2-PD centroids onto the corresponding AF2 structure and applying the same agglomerative clustering with a 10 Å threshold: clusters containing centroids from both datasets were flagged as ligand-associated; those with only predicted centroids were treated as ligand-free. This step identified 1,787 ligand-associated AF2-PD pockets distributed across 533 clusters (Figure 2b). Removing these 533 clusters left 1,332 clusters composed exclusively of ligand-free pockets. Finally, to maximize the coverage of the globular human pocketome, we shortlisted the 100 clusters with the highest number of pockets.

**Figure 2.**
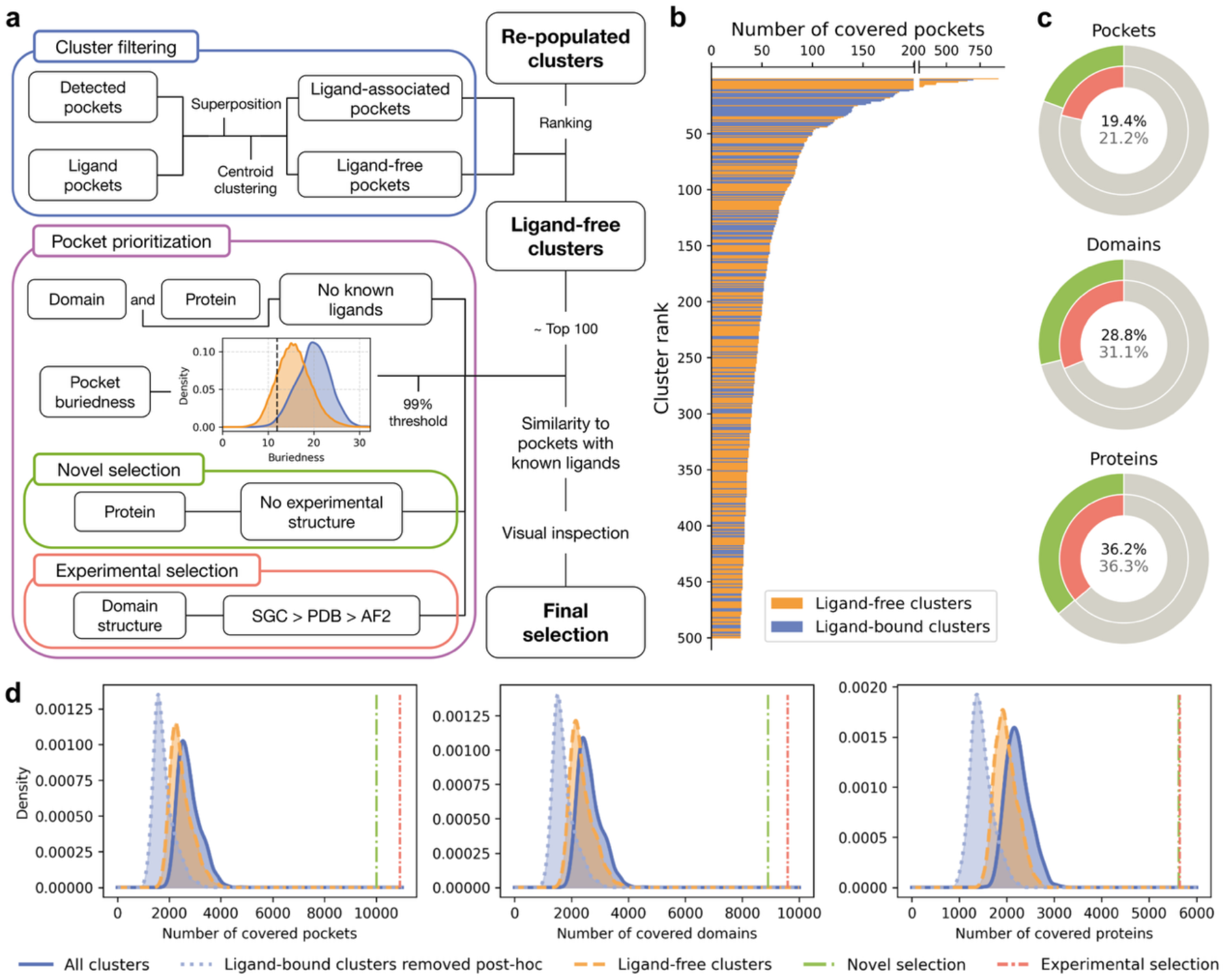
Obtaining the final representative pockets. **(a)** Schematic of the cluster filtering and pocket selection procedure. Clusters are ranked by number of descriptors; those containing at least one ligand-associated predicted pocket are removed. From the remaining ligand-free clusters, the 100 most populated are selected. Pockets from proteins or domains with known-ligand associations and pockets with buriedness < 11 are excluded. Two pockets per cluster with the lowest Mahalanobis distance to the ligand-associated pocket distribution are selected and visually assessed to obtain the final 200 representative pockets. **(b)** Number of descriptors for the 500 most populated clusters; clusters with at least one ligand-associated pocket (orange) and ligand-free clusters (blue). **(c)** Percentage of pockets, domains and proteins covered by the 100 selected clusters relative to the total number of pockets, domains and proteins for which a PocketVec descriptor was generated, for the novel (green) and experimental (red) selection strategies. Values are shown in black and gray for each selection respectively. **(d)** Distributions of the number of pockets (left), domains (middle), and proteins (right) covered by 100 randomly selected clusters (1,000 repetitions) from all clusters (dark blue, solid), after removing ligand-associated clusters from the previous selection (light blue, dotted), or from ligand-free clusters only (orange, dashed). Vertical dash-dot lines indicate the coverage of the novel (green) and experimental (red) selection strategies.

Within each cluster, additional filters were applied to enrich for pocket quality. Pockets from Pfam domains or proteins that harbor any known-ligand pocket elsewhere were excluded to avoid confounding effects in downstream experimental validation—since it would be difficult to disentangle whether observed binding signals arise from the pocket of interest or from a known ligand-binding site elsewhere in the same protein. Pockets with a buriedness below 11 were also removed; this threshold, derived from the distribution of ligand-associated AF2-PD pockets (Supplementary Figure 4), ensures that candidate sites have sufficient cavity depth for small-molecule binding.

From the 100 selected clusters, we picked two representative pockets per cluster from different domains, yielding a final set of 200 pockets for experimental characterization (Figure 2a). We decided to select two representatives per cluster to have some buffer against experimental failure and to increase the likelihood that at least one domain per cluster will yield measurable binding signals in AS-MS and DEL assays. The final set of 200 pockets was agreed with the SGC - Target 2035 team as a feasible number compatible with the preliminary pilot-phase throughput of the experimental pipeline.

To rank the filtered pockets within each cluster, we first examined five candidate quality metrics: the best, average, and worst docking scores of the 128 lead-like molecules used to derive the PocketVec descriptor, the P2Rank score, and buriedness. Comparing their distributions between ligand-associated pockets (those with an experimentally determined bound ligand) and *de novo* detected pockets, we found that best docking score, buriedness, and P2Rank score discriminated most clearly between the two groups (Supplementary Figure 4), and that none of the three was redundant with the others, so each contributes independent information (Spearman correlations of -0.46, -0.38 and 0.54 for docking score – P2Rank, docking score - buriedness and P2Rank - buriedness, respectively).

We then used these three metrics to define a reference distribution: the set of ligand-associated AF2-PD pockets, represented as a cloud of points in the three-dimensional metric space. For every candidate pocket we computed the Mahalanobis distance to this distribution — that is, its distance from the centre of the reference cloud, expressed in units of the cloud’s own spread and corrected for the covariance between the three metrics. Unlike a Euclidean distance, this makes the three metrics comparable despite their different scales and avoids double-counting the information they share. A low Mahalanobis distance therefore indicates a pocket whose combination of dockability, cavity depth, and predicted ligandability closely resembles that of experimentally validated binding sites, and which is consequently more likely to yield measurable binding. Candidates were ranked by this distance, and the two highest-ranked pockets per cluster were retained.

To handle the trade-off between scientific novelty and experimental tractability, we developed two complementary prioritization strategies applied after the Mahalanobis ranking. The novelty-focused strategy restricts candidates to proteins without experimentally determined structures in the PDB, prioritizing the structurally unexplored portion of the proteome. The experimental-focused strategy retains all candidates but stratifies them by structural availability—prioritizing domains whose structures are being (or have been) crystallized by the SGC, followed by domains with PDB structures, and finally domains represented only by AF2 models—before applying the Mahalanobis ranking within each tier. In both strategies, we assessed the final pocket candidates in three independent rounds of visual inspection in PyMOL ^34^, and those that did not exhibit pocket-like features (majority score of 1) were replaced by the next-ranked candidate from the same cluster, or the cluster was replaced by the next most populated qualifying cluster if no suitable replacement was available.

The novelty-focused strategy yielded 200 pockets from 197 domains (152 Pfam families) and 196 proteins. Additionally, the experimental-focused strategy yielded 200 pockets from 196 domains (163 Pfam families) and 185 proteins; of these, 98 belong to domains crystallized by the SGC, 87 to domains with PDB structures, and 15 to domains represented exclusively by AF2 models. Both strategies selected representatives from 100 clusters, of which 89 were shared, demonstrating that the coverage of the pocket landscape is primarily determined by the clustering and filtering steps rather than the final prioritization criterion. A 2D t-SNE projection of the full AF2-PD pocket space confirms that the 200 selected representatives are distributed uniformly throughout the landscape, covering all regions of the human pocketome rather than a specific chemical neighborhood (Figure 3).

**Figure 3.**
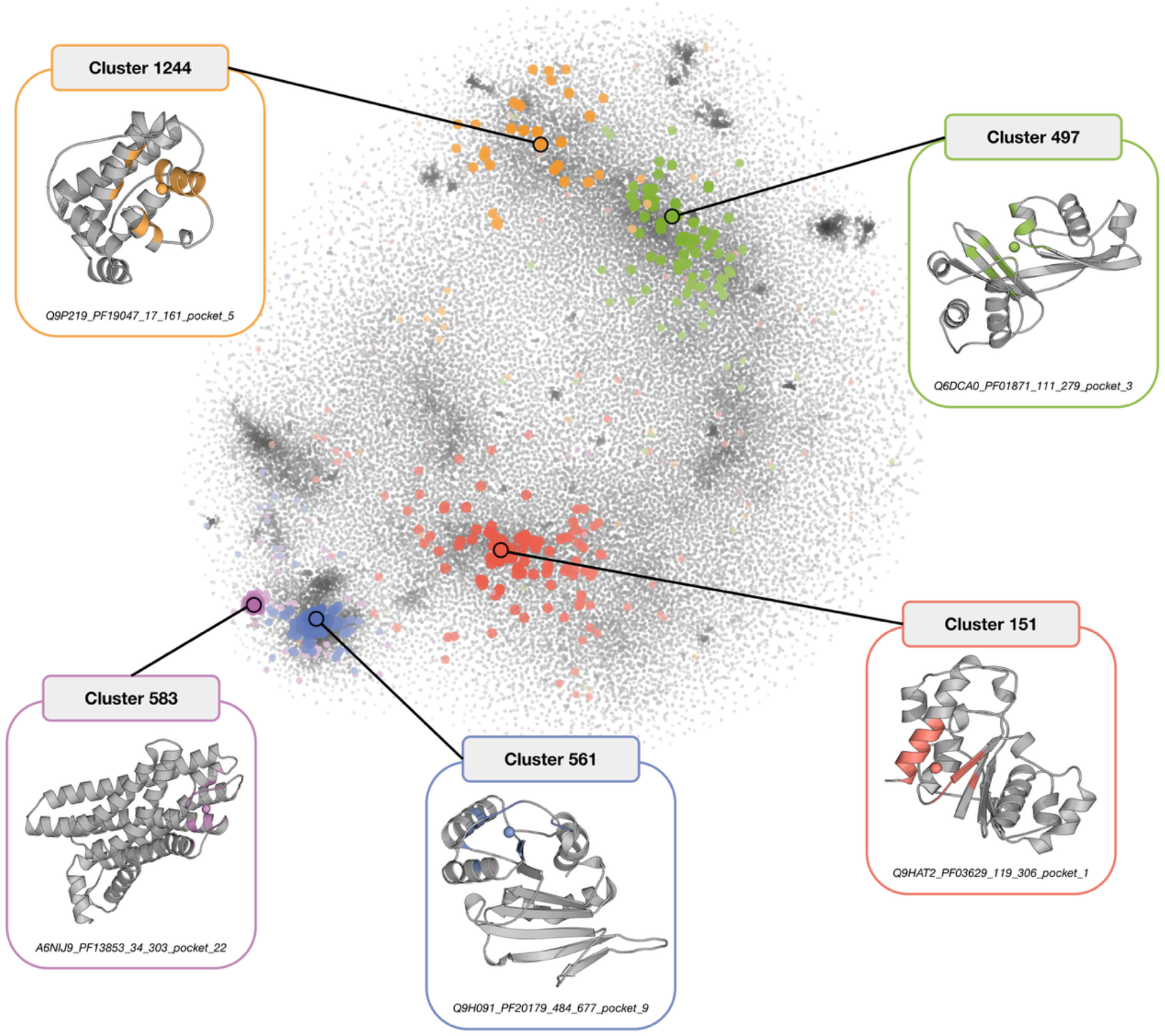
Overview of the generated pocket space. t-SNE projection of the AF2-PD PocketVec descriptors. Pockets belonging to five of the top 100 clusters are highlighted, while the rest are shown in gray. For each highlighted cluster, the structure and pocket centroid of one of the selected representatives, labeled with its identifier (Uniprot ID, Pfam domain, start and end positions and pocket number), is shown.

Collectively, the 100 selected clusters—and by extension the 200 final representatives— span ∼10,000–11,000 pockets (∼19–21% of the AF2-PD dataset), ∼8,900–9,600 domains (∼29–31%), and ∼5,600 proteins (∼36% of the human proteome with annotated pockets), for the novelty and experimental-focused strategies, respectively (Figure 2c). This leverage ratio—200 representatives covering information relevant to ∼5,600 proteins—is a direct consequence of the hierarchical similarity structure of the pocket space and demonstrates the power of pocket-based clustering as a diversity-maximizing target selection strategy. To contextualize these numbers, as stated above, we identified 1,787 AF2-PD pockets in 533 clusters with a co-crystallized ligand which, we extend the information to all pockets within these clusters, are representative of 18,570 pockets in 15,032 domains and 10,162 proteins.

In addition, to assess the added value of our selection scheme, we implemented three random baselines that profit from the clustered pocket space. In particular: (i) we selected 100 clusters at random from all 1,865 clusters; (ii) we applied the ligand-associated cluster filter after random selection, which discards any ligand-associated clusters and leaves a smaller set; and (iii) we applied the filter before random selection, i.e. selecting 100 clusters at random from the 1,332 ligand-free clusters, which is the baseline most comparable to our strategies (Figure 2d). All three baselines covered substantially fewer pockets, domains, and proteins than either of our strategies. Random selection from all clusters gave the highest random coverage (2,776 ± 432 pockets, 2,643 ± 410 domains, and 2,240 ± 252 proteins), followed closely by random selection from ligand-free clusters (2,464 ± 411, 2,357 ± 392, and 1,991 ± 233, respectively). In contrast, the novelty-focused and experimental-focused strategies covered 10,000 and 10,903 pockets, 8,891 and 9,581 domains, and 5,625 and 5,644 proteins, respectively, which represents a 2.8-to 4.4-fold increase over random selection from ligand-free clusters. Filtering after random selection was the least efficient option: on average, 28 of the 100 randomly selected clusters were ligand-associated and were discarded, reducing coverage by ∼1,000 pockets, domains, and proteins relative to selection from all clusters.

Overall, the coverage achieved by our strategies cannot be replicated by random cluster selection, even when ligand-associated clusters are excluded beforehand, indicating that the prioritization step is key to maximizing the fraction of pocket space represented. Moreover, the high proportion of ligand-associated clusters in random selections underscores the need to remove them before selection to obtain clusters that are both broadly representative and enriched in novel binding sites.

### Concluding Remarks

The generation of large, diverse protein–ligand datasets is a prerequisite for realizing the transformative potential of AI-based drug discovery. Target 2035 has set out an ambitious path to building such datasets^21^, but its success depends on selecting protein targets that collectively span the full chemical diversity of the human druggable proteome. Standard approaches based on sequence or fold similarity are not ideally suited to this task: binding-site similarity—not overall structural similarity—is the relevant criterion for predicting shared ligand-binding behavior.

The pocket-based framework presented here directly addresses this challenge. By applying PocketVec descriptors ^14^ to an updated and expanded map of more than 51,000 predicted binding sites across over 15,000 human proteins, and by clustering these sites at two levels of resolution—within Pfam families and across the entire proteome— we obtained a compact, non-redundant representation of the druggable pocket landscape. Selecting the 100 most populated chemically distinct, ligand-free clusters and choosing two high-quality candidate pockets per cluster yields 200 proteins whose experimental characterization will provide binding information relevant to approximately 36% of the human proteome—a leverage ratio that would be unachievable by random selection.

Two complementary prioritization strategies make the framework adaptable to different experimental priorities. When the primary goal is scientific novelty, the novelty-focused strategy restricts selection to structurally unexplored proteins. When experimental tractability is paramount, the experimental-focused strategy gives preference to proteins with available structures and active crystallization efforts by the SGC. Critically, both strategies achieve nearly identical coverage of the pocket landscape, confirming that broad chemical diversity is an intrinsic property of the clustering solution and is not sensitive to the final prioritization logic. Importantly, both strategies covered 2.8-to 4.4-fold more pockets, domains, and proteins than random selection of ligand-free clusters, indicating that this coverage is a direct result of our prioritization step rather than of chance.

We now look forward to the protein–small-molecule binding datasets that will be generated by our collaborators in the Target 2035 initiative. These data will allow us to evaluate the in vitro performance of the selected representative pockets, test the central prediction of our framework—that similar pockets will exhibit similar chemotype binding profiles in AS-MS and DEL assays—and iteratively refine the selection strategy for subsequent rounds of screening. Beyond Target 2035, the pocket-clustering approach is broadly applicable to any large-scale chemical biology effort where resource constraints require principled, diversity-maximizing target selection.

## Methods

### Generating PocketVec descriptors

We generated PocketVec descriptors for all druggable pockets in the human proteome following the methodology described in^14^, with several modifications to expand proteome coverage. We retrieved all reviewed human protein sequences from UniProt ^26^ and annotated domains using Pfam^27^ via InterPro^28^, including entries classified as “domain” or “family”. We structurally annotated the domains using experimental structures from the PDB^30^ and predicted models from the AlphaFold Protein Structure Database^31^. Relative to the original pipeline, we additionally incorporated mmCIF-format PDB structures^29^ and AF2 models for proteins exceeding 2,700 amino acids.

We identified candidate druggable pockets using a ligand-based approach (LIG), which defines pockets from co-crystallized ligands meeting the following criteria: PDBSUM-annotated, non-proteinogenic, containing more than 6 carbon atoms, and with solvent accessibility ≤ 0.4 or buriedness ≥ 15. We identified de novo pockets using Fpocket^10^ and P2Rank^11^, retaining the top two pockets per domain, provided that all residues within 8 Å of the centroid had pLDDT ≥ 70 for AF2 models. We prepared structures with PDBFixer^32^. We generated PocketVec descriptors by docking 128 lead-like molecules with rDock^35^ rigid docking and ranking the scores, and discarded descriptors with more than 80 outlier scores. We calibrated similarity thresholds by benchmarking on ProSPECCTs^13^, as described in the original PocketVec publication^12^ (Supplementary Figures 1 and 2).

### Defining pockets in Pfam domains

For each Pfam family, we selected a reference structure and superimposed all remaining domain structures, together with their pocket centroids, onto it using TM-align^33^. We computed pairwise Euclidean distances between centroids within the same family using SciPy^36^ and clustered the centroids using complete-linkage agglomerative hierarchical clustering with a 10 Å threshold, which we determined from the bimodal distribution of centroid-to-centroid distances (Figure 1b). We treated each resulting cluster as a single shared Pfam-level pocket. We validated these pocket definitions by comparing intra-pocket, inter-pocket, and inter-domain cosine distance distributions using two-sample KS tests (Figs. 1c,d).

### Creating and clustering the reduced dataset

For each Pfam-level pocket, we selected the centroid with the minimum mean cosine distance to all other members as the representative descriptor. We validated the resulting reduced dataset by confirming that its inter-descriptor distance distribution closely matched that of the full AF2-PD dataset (Figure 1e).

We clustered the reduced AF2-PD dataset using complete-linkage agglomerative hierarchical clustering with a cosine distance threshold of 0.22, as implemented in Scikit-learn^37^. We evaluated cluster quality by comparing intra- and inter-cluster cosine distance distributions using two-sample KS tests (Figure 1f). We then re-assigned the non-representative Pfam-level pockets to the cluster of their corresponding representative (Figure 1g).

### Selecting the pockets of interest

We excluded clusters containing at least one ligand-associated predicted pocket. We determined ligand association by superimposing AF2-LIG and AF2-PD centroids onto the corresponding AF2 structure using TM-align and applying agglomerative clustering with a 10 Å threshold; we flagged clusters containing centroids from both datasets as ligand-associated. We retained the 100 most populated ligand-free clusters. Within each cluster, we excluded pockets if their Pfam domain or protein contained any known-ligand pocket, or if their buriedness was below 11 (Supplementary Figure 4). In the novelty-focused strategy, we additionally excluded pockets from proteins with PDB structures. In the experimental-focused strategy, we stratified pockets by structural availability (SGC-active > PDB > AF2-only).

For the experimental-focused strategy, we identified SGC-associated pockets using a curated list of SGC constructs with UniProt identifiers and residue boundaries. After removing complexes and non-human entries, we retained domains whose boundaries were contained within a synthesized SGC construct, which yielded 6,897 SGC-associated pockets across 3,930 domains and 2,417 proteins.

### Final protein selection

From the filtered candidates in each of the 100 selected clusters, we selected two representatives per cluster by ranking candidates according to their Mahalanobis distance^38^ to the multivariate distribution of ligand-associated pockets, defined by best docking score, buriedness, and P2Rank score (Supplementary Figure 4). In the experimental-focused strategy, we first stratified candidates by structural tier (SGC, PDB, AF2) and then ranked them by Mahalanobis distance within each tier. We assessed all final candidates through three independent rounds of visual inspection in PyMOL^34^. We replaced non-pocket-like candidates (majority score of 1) with the next-ranked candidate from the same cluster or, if no suitable replacement was available, replaced the cluster with the next most populated qualifying cluster.

## Data and Code availability

All the data used in this study, as well as the code developed is publicly available at https://github.com/sbnb-irb/Target2035/. Detailed documentation describing the workflow, required input data, and repository structure is also provided.

## Acknowledgment

PA acknowledges the support of the Spanish Ministerio de Ciencia, Innovación y Universidades (PID2023-152296OB-I00) and the European Commission - Horizon Europe Project CLARITY (101137201). We also acknowledge institutional funding from the Spanish Ministry of Science and Innovation through the Centres of Excellence Severo Ochoa Award, and from the CERCA Programme / Generalitat de Catalunya.

## Ethics Declarations

The authors declare no conflict of interest.

This project has been partially funded by the European Union. However, views and opinions expressed are however those of the author(s) only and do not necessarily reflect those of the European Union or the Health and Digital Executive Agency. Neither the European Union nor the granting authority can be held responsible for them.

## Supplementary Figures

**Supplementary Figure 1.**
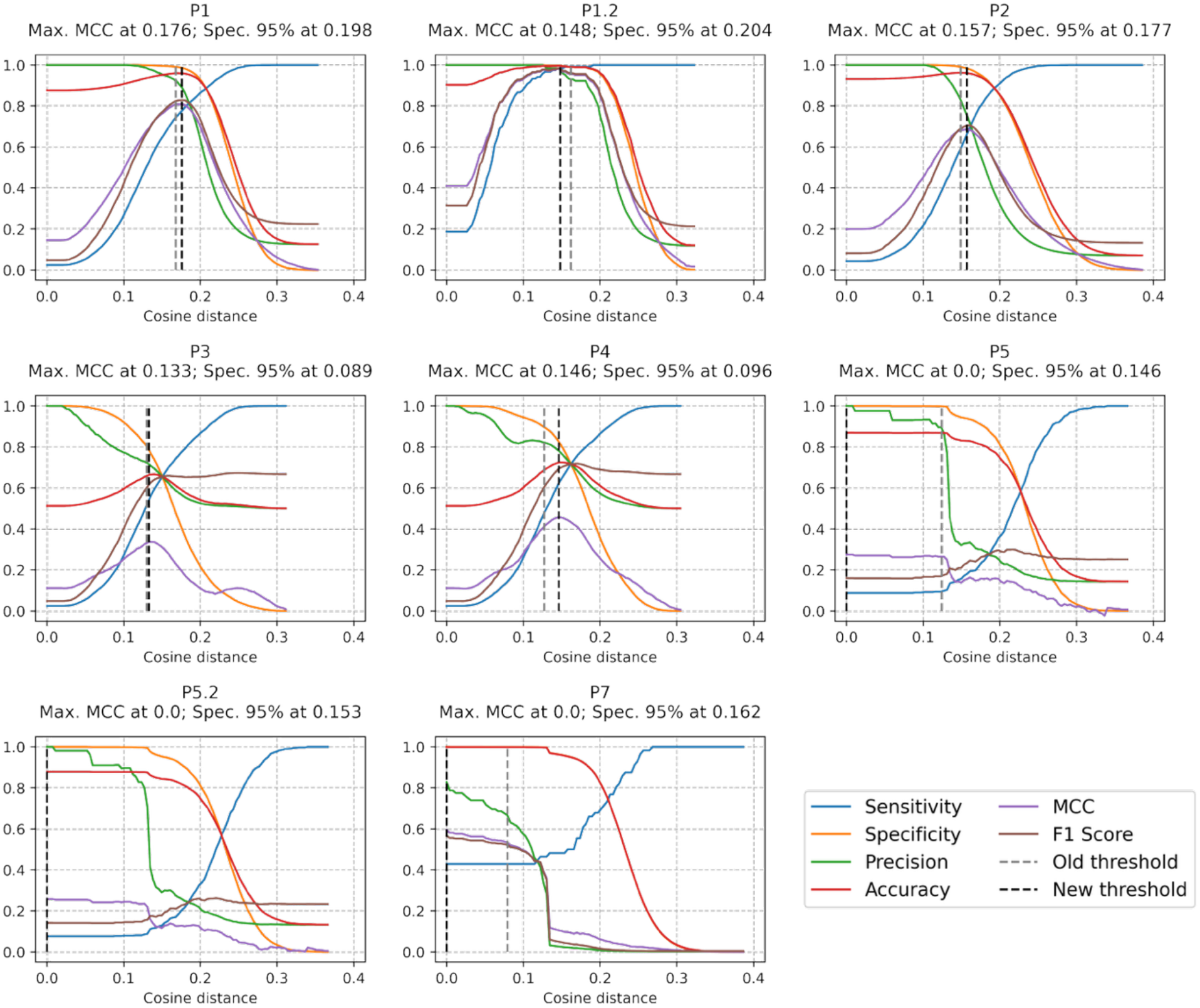
Benchmarking of updated PocketVec descriptors on the ProSPECCTs dataset for PDB structures. AUROC values across all seven ProSPECCTs datasets. Dataset P1 was used to calibrate the cosine distance threshold for PDB-derived structures (0.18 in the updated pipeline vs. 0.17 in the original publication).

**Supplementary Figure 2.**
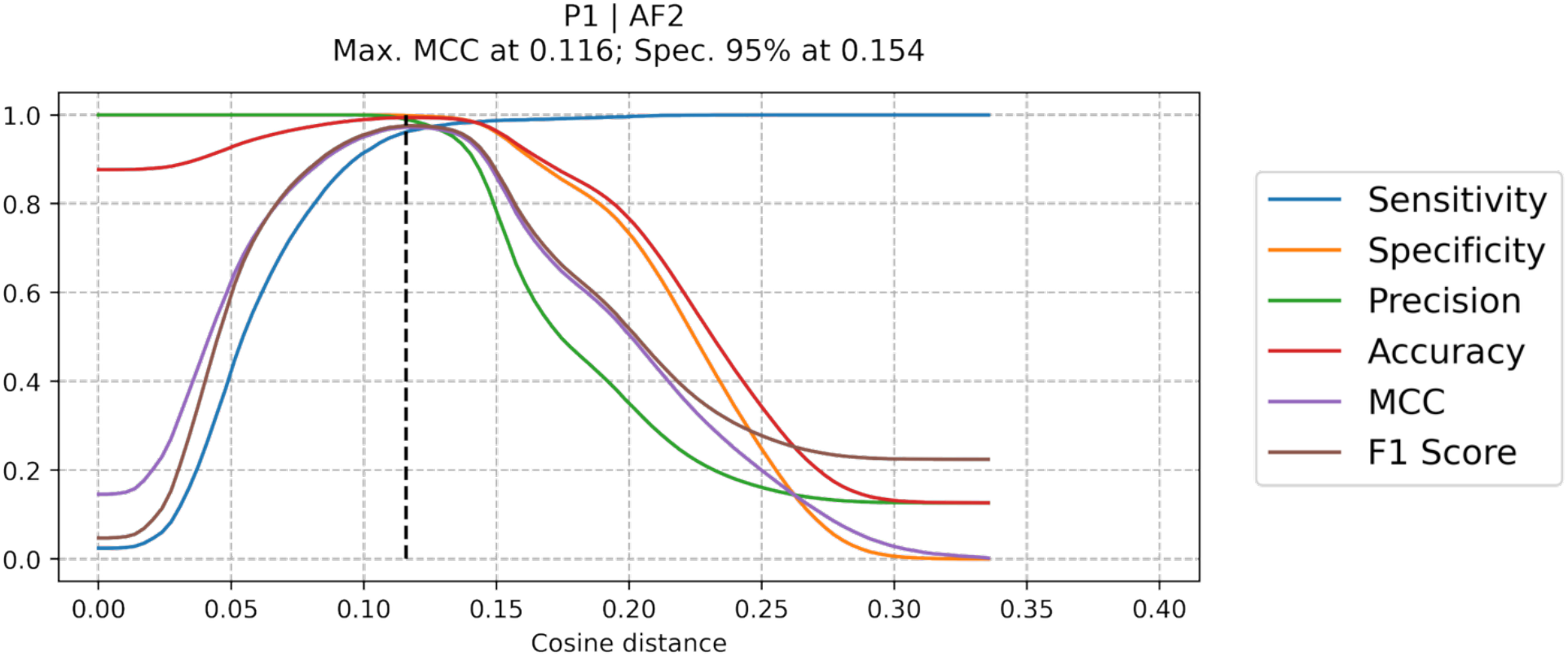
Benchmarking of updated PocketVec descriptors on the ProSPECCTs dataset for AF2 structures. Benchmarking was restricted to dataset P1. The cosine distance threshold for AF2-derived structures was set at 0.16.

**Supplementary Figure 3.**
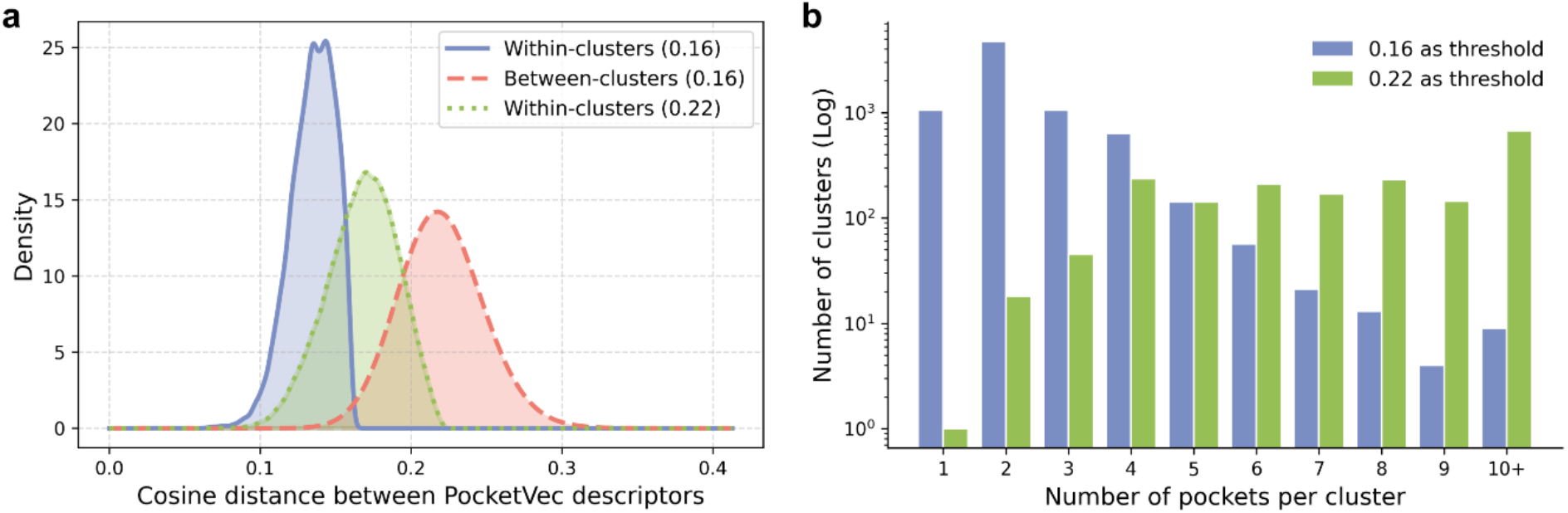
Separation between similar and dissimilar pockets and number of pockets per cluster. **(a)** Intra-cluster cosine distance distributions for the reduced AF2-PD dataset clustered using a 0.16 (blue) and 0.22 (green) distance threshold. Inter-cluster cosine distance distribution is shown for the 0.16 clustering only (red), as it is nearly identical at 0.22. **(b)** Bar plot of the number of clusters generated with a threshold cosine distance of 0.16 (blue) and 0.22 (green) by number of included pockets

**Supplementary Figure 4.**
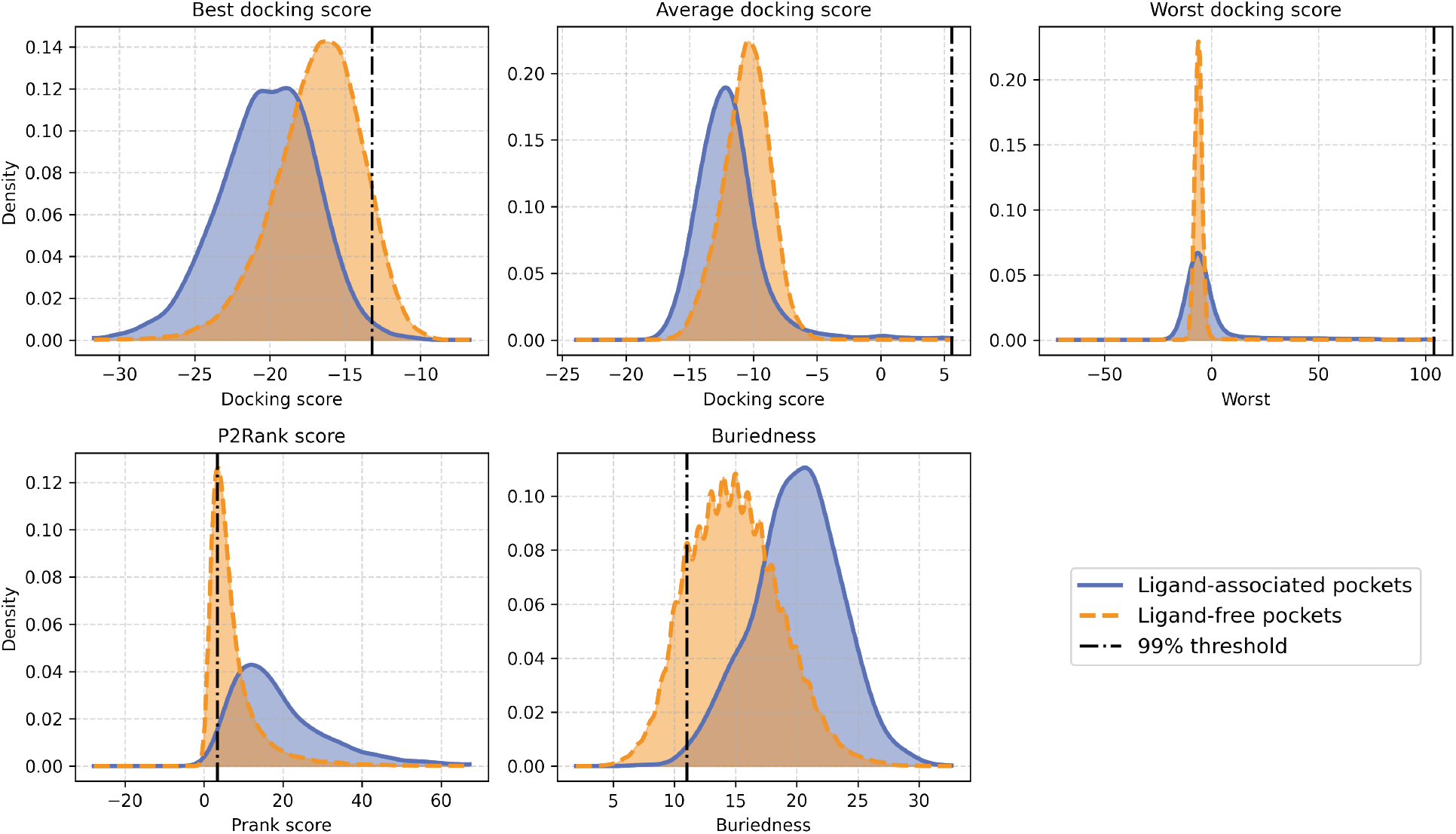
Pocket quality metrics for ligand-associated and ligand-free AF2-PD pockets. Distributions of the five metrics used for quality assessment: best, average, and worst docking scores; P2Rank score; and buriedness., shown separately for ligand-associated (blue, solid) and ligand-free (orange, dashed) predicted pockets. The 99% threshold is indicated for all metrics. The three metrics selected for the Mahalanobis distance calculation (best docking score, buriedness, P2Rank score) were chosen based on correlation analysis to maximize complementary information.

